# Chemogenetic inhibition of amygdala to ventrolateral prefrontal cortex communication selectively impacts contingent learning

**DOI:** 10.1101/2025.06.26.661815

**Authors:** J. Megan Fredericks, Catherine Elorette, Jared E. Boyce, Peter H. Rudebeck

**Affiliations:** Nash Family Department of Neuroscience, Lipschultz Center for Cognitive Neuroscience, and Friedman Brain Institute, Icahn School of Medicine at Mount Sinai, One Gustave L. Levy Place, New York, NY, 10029, USA

**Author notes:** Department of Neurobiology, University of Pittsburgh, Pittsburgh, PA, USA. Medical Scientist Training Program, University of Wisconsin-Madison, Madison, WI, USA. **CORRESPONDENCE SHOULD BE ADDRESSED TO**: Dr. Peter H. Rudebeck Icahn School of Medicine at Mount Sinai One Gustave L. Levy Place New York, NY, 10029, USA.

**Keywords:** Amygdala, ventrolateral prefrontal cortex, learning, choices, valuation, probability, reward

## Abstract

Contingent learning, the process by which specific courses of action become associated with subsequent outcomes, is dependent on the amygdala and ventrolateral prefrontal cortex (vlPFC). The amygdala and vlPFC are bidirectionally connected but it is unclear what the contribution of individual feedforward and feedback pathways is to contingent learning. Here we tested the role of amygdala projections to vlPFC in mediating two key components of contingent learning: signaling the outcome (reward/no reward) that follows a choice and maintaining representation of the choice that was made prior to outcome delivery. To test for these two aspects of contingent learning, we trained macaques to perform a probabilistic reward learning task where for separate stimulus pairs reward was either delivered immediately or after a trace interval. Inhibiting vlPFC-projecting amygdala neurons impacted contingent learning irrespective of whether there was a trace interval or not, and this effect was primarily driven by maladaptive learning on unrewarded trials. Notably, deficits in contingent learning caused by manipulating activity in the amygdala-vlPFC pathway were distinct from impairments in motivation and the ability to update the value of specific rewards in a reinforcer devaluation task. Thus, vlPFC-projecting amygdala neurons appear to play a specific role in contingent learning through signaling the outcomes of a choice, but not in maintaining a memory of the prior choice.

**SIGNIFICANCE STATEMENT:** Impairments in learning which stimuli or actions are associated with temporally proximate rewards or punishments is characteristic of a host of psychiatric and neurological disorders. Determining the neural circuits that support this type of contingent learning is therefore a key question in neuroscience. Here we show that that during contingent stimulus-reward learning, projections from amygdala to ventrolateral prefrontal cortex (vlPFC) in macaques are essential for signaling reward delivery or omission. These projections appear to be less involved in maintaining a memory of the prior choice that led to the reward or punishment that is subsequently experienced or for updating the value of the received rewards. Thus, amygdala-vlPFC connections make a specific contribution to signaling reward feedback during learning.

## INTRODUCTION

Quickly and adaptively learning which course of action in our environment will be most likely to lead to rewards or punishments is critical for survival. This type of learning, known as contingent learning, relies on linking together the course of action that was taken with the reward or punishment that is subsequently experienced. Recent studies in macaque monkeys using various approaches including excitotoxic lesions^1,2^, focused ultra-sound^3^, neuroimaging^3,4^, and neurophysiology^5–8^ suggest the importance of ventrolateral prefrontal cortex (vlPFC), centering on Walker’s area 12, and amygdala for contingent learning in probabilistic settings. Given this pattern of effects, and because vlPFC and amygdala send and receive direct monosynaptic projections^9,10^, these two areas may form part of a circuit specialized for contingent learning in primates^11^.

What is not presently clear, however, is whether the feedforward projections from amygdala to vlPFC play a role in contingent learning and what specific aspect of contingent learning they might contribute to. One possibility is that amygdala inputs to the vlPFC are required to signal the presence or absence of a biologically salient event, such as a reward or punishment that follows a particular choice so that stimulus-reward associations can be iteratively updated. This would fit with findings that amygdala lesions slow contingent learning^1,12,13^, and reduce the representations of expected rewards in frontal cortex^12,13^, and that single neurons in amygdala encode the presence or absence of rewards and punishments following a choice^14^.

Another possibility is that projections from amygdala to vlPFC are required to solve the temporal credit assignment problem, a key component of contingent learning. Specifically, contingent learning relies on there being a temporal contingency between the stimulus or action that is selected and the reward or punishment that is subsequently experienced^15,16^, i.e. the outcome has to be “credited” to the prior choice. Consequently, learning is slowed if the period of time between a choice and the outcome of that choice, known as a trace interval, increases^16^. Unlike a delay period, in a trace interval no information is available about the preceding choice. This means that a trace memory about the stimulus selected or action taken must be maintained during the length of the interval to effectively associate it with the outcome that follows. This influence of trace intervals on learning is consistently seen across humans^17,18^, apes^19^, monkeys^20^, rodents^21^ and other species^16,22^ demonstrating that this process is evolutionarily conserved. In macaques, prior work has found that amygdala neurons exhibit sustained encoding during trace intervals^14,23^ and also track reward timing^24^, indicating this area could be required to maintain a memory trace of the stimulus or action that will become associated with the subsequent outcome.

To assess these two possible roles for the feedforward projections from amygdala to vlPFC in contingent learning, we tested animals on a probabilistic-reward learning task with two conditions: one where a trace period was present after the animal selected a stimulus, and one without a trace period. We used a dual-viral chemogenetic approach to selectively express inhibitory DREADD (Designer Receptors Exclusively Activated by Designer Drugs) receptors in only those amygdala neurons projecting to vlPFC. If the amygdala-vlPFC pathway is essential for signaling the outcome of a choice (i.e. reward or no reward), then inhibiting these projections should impair contingent learning irrespective of whether a trace interval is present or not. Alternatively, if amygdala-vlPFC projections are maintaining a memory of the choice across the trace interval, then their inhibition should only impair contingent learning when a trace interval is present, and a memory of the choice is necessary to learn stimulus-reward associations. We observed a similar deficit in learning across both the trace and no trace conditions after inhibition of vlPFC-projecting amygdala neurons. Thus, our data indicate that amygdala-vlPFC projections contribute to contingent learning by signaling the presence or absence of reward as opposed to maintaining a memory of the prior choice.

## RESULTS

To establish the role of amygdala to vlPFC projections in contingent learning, we trained three macaque monkeys (Cy, Wo, and He) to perform a probabilistic stimulus-reward learning task for food rewards (**Figure 1A**). In each 200-trial session, subjects had to discriminate two pairs of visual stimuli on a touch sensitive monitor. A trial began when subjects touched a green ‘lever’ stimulus centrally presented on the screen. A pair of stimuli were then presented to the left and right of the screen. In each pair, choosing one of the stimuli led to reward 80% of the time whereas choosing the other led to reward 20% of the time. For one pair of stimuli, the outcome (reward or no reward) was delivered immediately, while for the other pair there was a 1 second trace interval between the choice and the outcome delivery. During this interval the chosen stimulus was removed from the screen and animals had to maintain a “trace” of their prior stimulus choice. The stimulus pair presented on each trial was pseudo-randomly selected, each pair was presented for a total of 100 trials in each session, and the left-right position of each stimulus in a pair was counter balanced. The stimuli in each pair were novel at the start of each session and the assignment of stimuli to each level of reward probability and pair was random.

**Figure 1:**
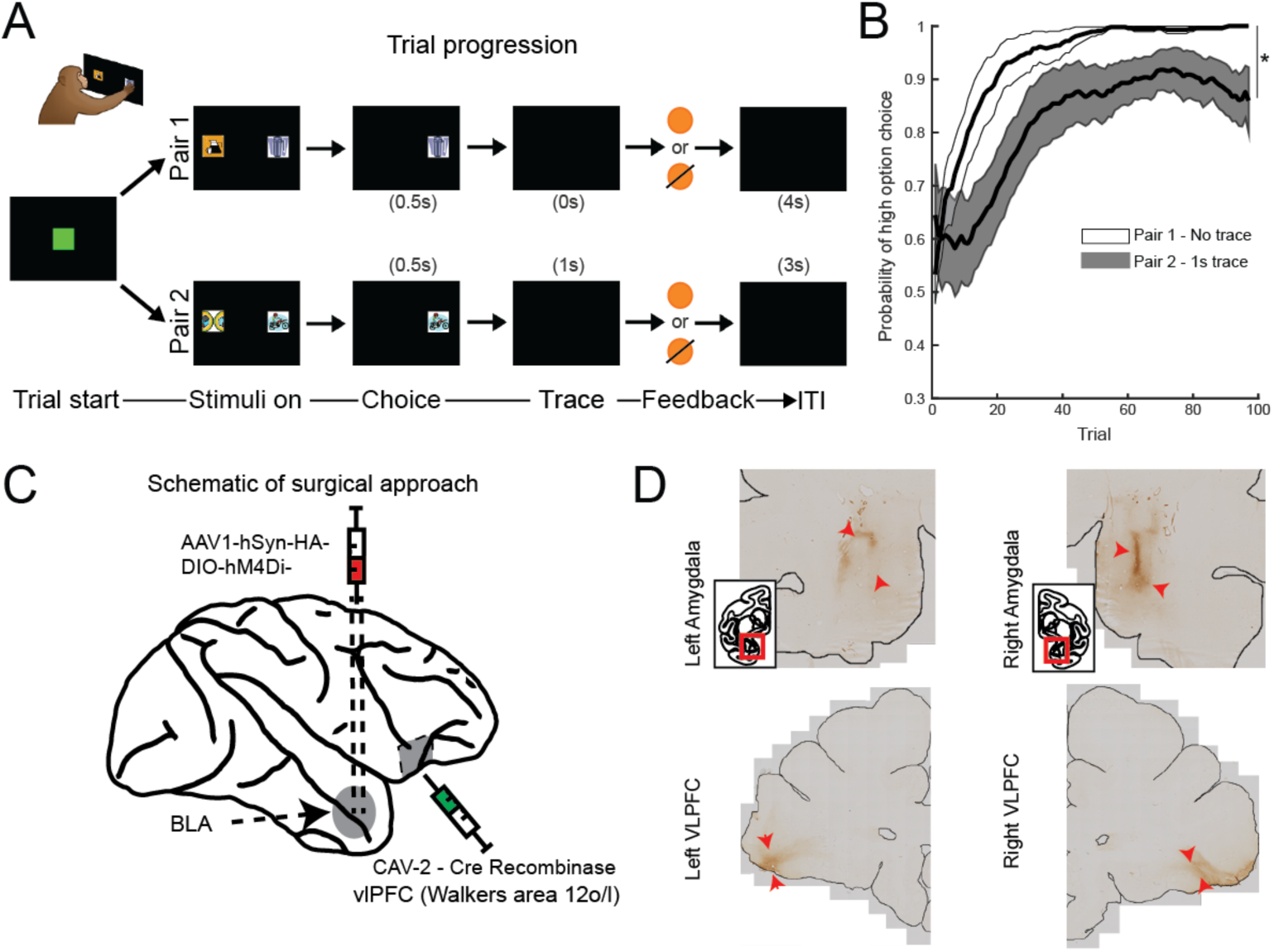
Behavioral task, subject’s performance, surgical approach, and immunohistology. A) Trial sequence of the two-pair probabilistic learning task. Each trial started when subjects touched the green ‘lever’ stimulus. This led to the presentation of either pair 1 or 2. In each pair of stimuli one stimulus was associated with 0.8 probability of reward and the other with 0.2. Choosing one of the stimuli led to it being presented alone for a further 0.5 seconds. For pair 1, reward feedback was then delivered when the chosen stimulus was extinguished, whereas for pair 2 reward feedback was delivered after a 1 second trace interval. B) Subjects were slower to learn the option associated with the higher probability of reward when there was a 1 second trace interval between choice and reward feedback. * denotes p<0.05. C) Schematic of surgical approach. D) Light microscopy images of the left and right amygdala (top) and vlPFC (bottom) for immunohistologically stained sections against the HA-tag. Outlines for the sections are included to show the limits of the brain. Insets show location of section being shown relative to the rest of the brain. Red arrows point to stained areas.

To ensure that the temporal reward rate was the same for both pairs of stimuli, we adjusted the intertrial interval (ITI) such that the time from a choice to the start of the next trial was always 4 seconds. Extensive piloting of the task design showed that this reduced the delay-discounting effect, as performance was even more impaired when the ITI was not adjusted to maintain the temporal reward rate (**Supplementary Information, Figure S1**). To balance efficient and consistent learning with a clear effect of the trace interval on learning, we selected the 1s trace interval to compare to immediate reward feedback. Mirroring what has been found across many species^3–8^, the presence of the trace interval impacted how many trials it took subjects to discriminate the stimulus that was associated with the higher probability of reward (**Figure 1B**). Specifically, subjects were slower to learn when there was a 1 second trace interval between their choice and reward delivery (logistic regression, effect of pair, t=9.24, p<0.0001). Similarly, subjects took longer to choose a stimulus in the trace condition (choice response time, effect of stimulus pair, F(_1,5987_)=32.5, p<0.0001).

Next, we confirmed that the DREADD activating ligand, deschloroclozapine (DCZ),^25^ had no effect on subject’s performance prior to DREADD injection surgery. Here, all subjects were tested with doses of 0.1 and 0.3 mg/kg DCZ while they performed the dual pair probabilistic stimulus-reward learning task. 0.1 mg/kg DCZ had minimal impact on task performance compared to saline (logistic regression, effect of drug X^2^= 1.08, p > 0.05) while 0.3 mg/kg DCZ had a noticeable impact on animals learning behavior (logistic regression, effect of drug X^2^= 5.69, p < 0.01) as has been reported previously^26^. The higher dose of DCZ also impacted reaction times (ANOVA, effect of drug, F_1,35_ = 14.03, p < 0.01) in the task and impacted the time to press the green square to initiate trials (ANOVA, effect of drug, F_1,35_ = 5.78, p < 0.05), whereas 0.1 mg/kg did not impact either measure of performance (both comparisons, ANOVA, effect of drug, F_1,35_ < 2.34, p > 0.13). Thus, all subsequent testing proceeded with DCZ given at a concentration of 0.1 mg/kg.

In an aseptic neurosurgery, MRI-guided stereotactic injections of AAV1-DIO-hSyn-hM4Di-HA and CAV2-Cre^27^ were made bilaterally into the amygdala and vlPFC in each subject, respectively (**Figure 1C**). In amygdala, we targeted injections to the basal and accessory basal nuclei and in vlPFC, injections targeted Walker’s area 12 on the ventral surface of the frontal lobe. This intersectional virus approach meant that only neurons projecting from amygdala to vlPFC would express the inhibitory DREADD receptor. The expression of the DREADD receptors in basal amygdala was verified immuno-histologically (**Figure 1D**, top panels) and we confirmed the presence of labeled processes in vlPFC (**Figure 1D**, bottom panels). All postoperative testing with DCZ was conducted within 2 years of the date of surgery to ensure adequate DREADD receptor expression^28^.

Inhibiting the pathway from amygdala to vlPFC caused a deficit in learning that was apparent for both the trace and no trace conditions (logistic regression, effect of drug, t=-8.766, p<0.001, **Figure 2A**) although it is important to note that the difference in learning between the conditions was maintained (logistic regression, effect of pair, t=-9.24, p<0.00001). The deficit was, however, more pronounced for the no trace condition (logistic regression, drug by pair interaction, t=6.1, p<0.001; *post hoc* test, pair 1, effect of drug, t=10.91, p<0.0001). A similar decrement in performance was also seen for learning when a the trace interval was present (logistic regression, effect of drug, pair 2, t=-5.64, p<0.00001). The effect of inhibiting the pathway from amygdala to vlPFC was most apparent during the first 50 trials of the session for each pair (logistic regression, block by pair by injection interaction, t=3.88, p<0.0002) indicating that the effects were most prominent during the phase where subjects were learning the probabilities associated with the stimuli. Thus, while inhibiting the pathway from amygdala to vlPFC had a slightly greater effect on learning when reward was delivered immediately, it had a non-selective effect on learning, as it impacted learning in both the trace and no trace conditions. This suggests that rather than maintaining a memory trace of the chosen stimulus, amygdala-vlPFC projections contribute to contingent learning through signaling information about the outcome of a choice to guide subsequent choices.

**Figure 2:**
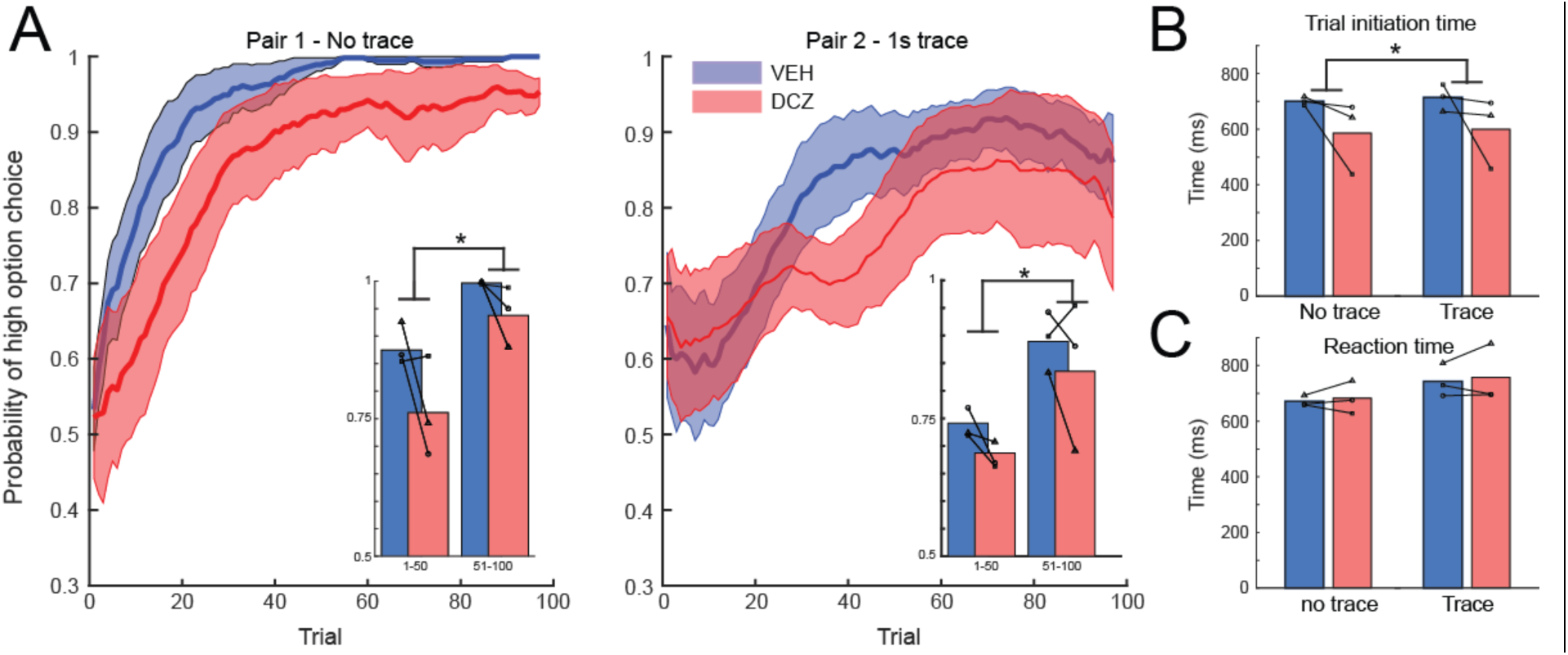
The effect of inhibiting amygdala to vlPFC projections on probabilistic learning. A) Mean (+/- S.E.M.) probability of choosing the stimulus associated with the high probability of reward for the no trace (left) and trace interval (right) pair of stimuli. Drug condition is signified by color; blue, vehicle (VEH) and red, deschloroclozapine (DCZ). Inset figures show the mean high option choice probability for the first (trials 1-50) and second (51-100) blocks of trials. B) Mean initiation and C) reaction times for the no trace and trace pairs of stimuli. In all plots, individual symbols connected by lines denote individual subject performance. * denotes p<0.05.

Inactivating the pathway from amygdala to vlPFC also had a differential effect on the time that it took subjects to initiate trials (trial initiation time, **Figure 2B**) and then choose either of the two stimuli (reaction time, **Figure 2C**). Specifically, administration of DCZ increased initiation time to press the ‘lever’ stimulus to start a trial for both trace and no trace conditions (ANOVA, effect of drug, F(_1,11,984_)=34.15, p<0.001; drug by pair interaction, F(_1,11,984_)=0, p>0.9). By comparison, there was no difference in reaction times to choose between the two stimuli in either condition when DCZ was administered (effect of drug F(_1,11,984_)=1.48, p>0.2; pair by drug interaction). We interpret the effect on initiation, but not reaction times as a result of subjects generally being less able to form contingent stimulus-reward associations as opposed to changes in motivation which would have affected both measures.

### The influence of feedback on subsequent choices

To further pinpoint the specific role of the projections from amygdala to vlPFC we next investigated how choices and their contingent feedback influenced future behavior (**Figure 3A**). First, we looked at subjects’ behavior under control conditions, when vehicle injection was administered and thus communication between amygdala and vlPFC was intact. As learning progressed during a session, subjects were more likely to ‘stay’, or continue choosing, a previously rewarded option (win-stay behavior, effect of block, t=4.53, p<0.0001). This effect was smaller for the no trace condition, likely as subjects’ learning asymptotes sooner for this pair (**Figure 3A**, win-stay behavior, block by pair interaction, t=-2.99, p<0.005). A slightly different pattern emerged for behavior following unrewarded trials. Here subjects were less likely to shift their choices after not receiving a reward for the no trace condition, compared to the trace condition (**Figure 3B**, lose-shift behavior, effect of pair, t=-3.56, p<0.0005). This effect was most apparent in the later stages of each session for the no trace condition as subjects’ valuations of the two stimuli stabilized (lose-shift behavior, block by pair interaction, t=-3.56, p<0.0005).

**Figure 3:**
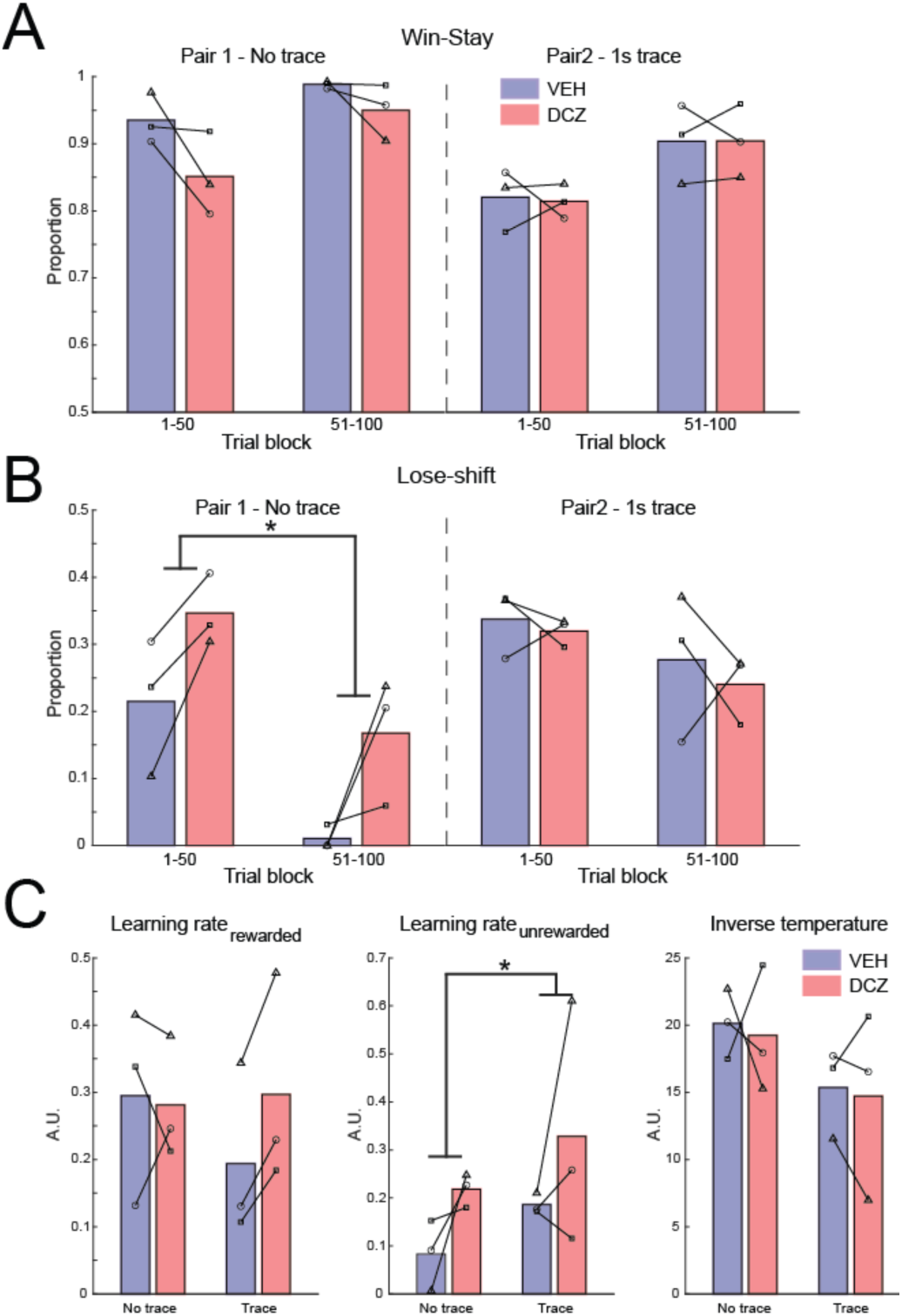
Influence of previous feedback on behavior and reinforcement learning models. A) Mean win-stay and B) lose-shift behaviors for each pair and drug condition during the first (trials 1-50) and second (trials 51-100) block of trials. C) Mean modeled free parameters, learning rate for rewarded, learning rate for unrewarded trials, as well as inverse temperature from a reinforcement learning model fitted to subjects’ choices. Drug condition is signified by color; blue, vehicle (VEH) and red, deschloroclozapine (DCZ). Individual symbols connected by lines denote individual subject performance. * denotes p<0.05.

Next, we looked at how inhibition of the neurons projecting from amygdala to vlPFC altered how feedback was used. When DCZ was administered, lose-shift behavior was increased in all subjects, but only when there was no trace present (**Figure 3b**, lose-shift behavior, pair by block by drug interaction, t=-3.75, p<0.0002; *posthoc* test, pair 1, effect of drug, t=-2.27, p<0.05). By contrast, while there was a slight decrease in the use of win-stay strategies after DCZ administration for the no trace condition, this effect failed to reach statistical significance (**Figure 3A**, win-stay behavior, effect of drug and interactions involving drug, t<1.5, p>0.1). Taken together, inhibition of amygdala-vlPFC projections caused a selective increase in switching behavior following unrewarded choices when feedback was delivered immediately.

This effect of unrewarded trials on subsequent behavior potentially indicates that inhibiting the amygdala-vlPFC pathway causes subjects to decrease their reward valuations of the current option to a greater degree after not receiving a reward. To further characterize this effect, we fitted a reinforcement learning model to animals’ choices. Based on the prior finding of differences between rewarded and unrewarded trials (**Figure 3A/B**), the model included separate learning rates for rewarded and unrewarded trials as well as an inverse temperature parameter (see **Materials** and **Methods**). The model was also fitted to choices for each pair of stimuli separately and the three free parameters were estimated using standard approaches. Inhibiting amygdala-vlPFC projections increased the learning rate for unrewarded trials irrespective of whether feedback was delivered after a trace interval or not (**Figure 3C**, unrewarded learning rate, ANOVA, F(_1,114_)=8.55, p<0.005). This effect was limited to unrewarded trials, as the learning rate for rewarded trials and the inverse temperature parameter were unaffected (ANOVA, F(_1,114_)<1, p>0.1). Taken together, these analyses show that inhibiting the pathway from amygdala to vlPFC selectively altered behavior following unrewarded trials.

### The influence of vlPFC-amygdala inhibition on updating stimulus-reward valuations

Finally, to confirm that the effects of amygdala-vlPFC pathway inhibition were specific to contingent learning, we tested whether inactivating these projections impacted animals’ ability to update the current value of specific rewards to guide their choices. This process is distinct from contingent learning as it requires updating the value of a specific outcome, in this case a food reward following a choice, not learning the association between a stimulus and the outcome that subsequently occurs. Here we used a reinforcer devaluation paradigm that has previously been shown to be sensitive to lesions of amygdala^29^. Subjects were trained to discriminate 60 pairs of stimuli (**Figure 4A**). Each pair contained one stimulus associated with the subsequent delivery of a specific food reward on every trial, while the other was unrewarded. Within the 60 pairs, 30 stimuli were associated with a peanut reward while the remaining 30 were associated with an M&M reward. The peanut or M&M reward designation for each stimulus was randomly assigned, set at the beginning of training, and was not subsequently changed, allowing subjects to learn and use stimulus-reward assignments across sessions.

**Figure 4:**
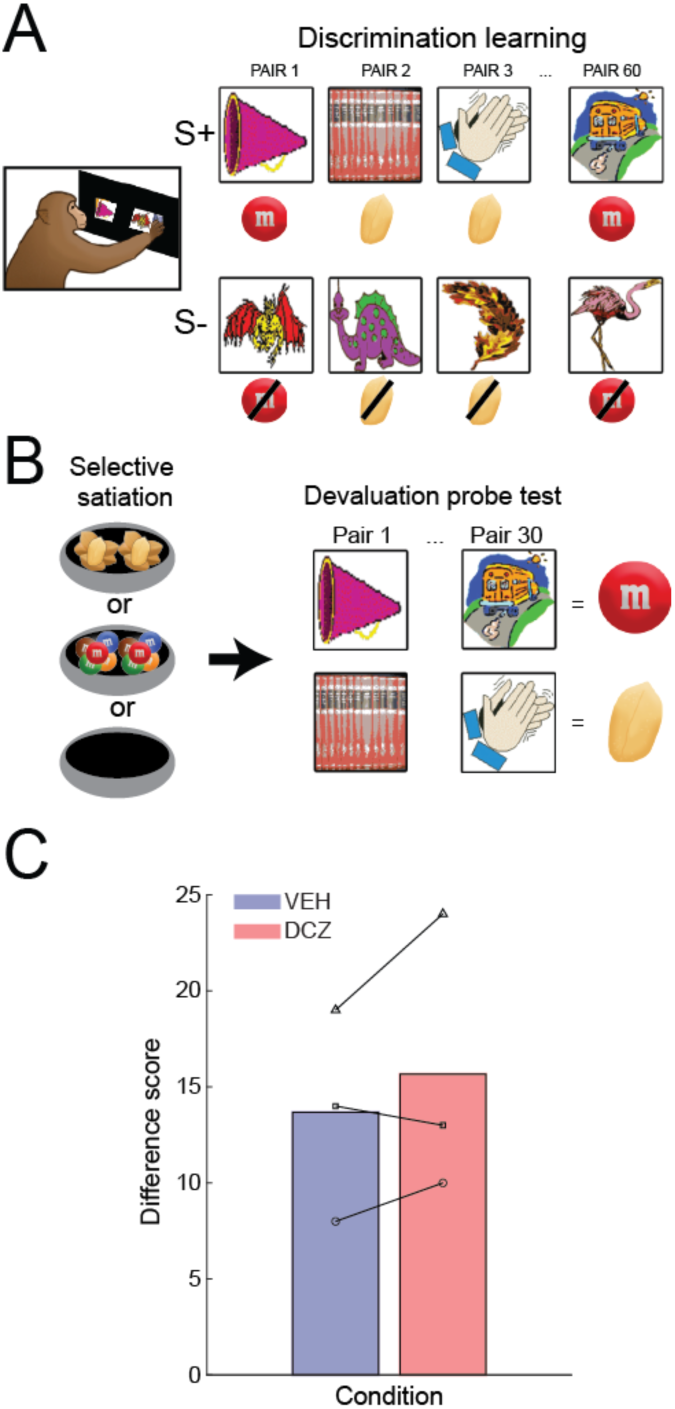
The effect of inhibiting the pathway from amygdala to vlPFC on a reinforcer devaluation task. A/B) Schematic of the discrimination learning (A) and devaluation probe sessions (B). C) Mean difference score when vehicle (blue, VEH) or DCZ (red) were administered. Individual symbols connected by lines denote individual subject performance.

Once subjects were successfully discriminating between the stimuli, critical test sessions involving novel pairs of reward-associated stimuli were performed (**Figure 4B**). This meant that on each trial a peanut associated stimulus was presented with an M&M associated stimulus. On set days, one of the reward options, either M&M or peanuts, was devalued using a selective satiation procedure (see **Materials** and **Methods**). Thus, by comparing behavior in sessions where a reward had been devalued to those where it had not, we could assess whether amygdala to vlPFC projections had a specific role in contingent learning or play a more general role in reward-guided behaviors that include updating the sensory-specific subjective value of rewards. Further, based on prior findings that amygdala is involved in updating the value of the outcomes as opposed to using updated valuations^30^, DCZ or saline were administered prior to the selective satiation procedure.

Inactivating the neurons projecting from amygdala to vlPFC did not alter subjects’ ability to update the value of specific rewards to guide their choices (**Figure 4C**). Inhibiting this pathway made subjects slightly more sensitive to changes in the value of specific rewards, mirroring prior observations of the effects of vlPFC lesions^2^, although this failed to reach statistical significance (saline versus DCZ, F_(1,5)_=0.14, p>0.5). Taken together, this pattern of results indicates that the neurons projecting from amygdala to vlPFC are essential for signaling reward outcome during contingent learning, not updating the subjective value of the reward that is received.

## Discussion

The present experiments reveal the selective role of the pathway linking amygdala to vlPFC in contingent stimulus-reward learning in macaques. Consistent with extensive prior work in humans and other animals^3–8^, macaques were slower to learn which option was most likely to lead to a reward when there was a trace interval between their choice and reward feedback (**Figure 1B**). We found that chemogenetically inhibiting the projections from amygdala to vlPFC impaired learning irrespective of the presence of a trace interval (**Figure 2**). This indicates that these projections are involved in signaling reward outcome during contingent learning as opposed to maintaining a representation of the choice across the trace interval. Further analyses revealed that the impact of pathway specific inhibition on performance was driven by an alteration in the way that subjects integrated unrewarded choices into their current valuations (**Figure 3**). Importantly, inhibiting the projections from amygdala to vlPFC was without effect on a reinforcer devaluation task that assessed subject’s ability to update the value of specific rewards that will follow a choice (**Figure 4**). This pattern of effects indicates that projections from amygdala to vlPFC are involved in contingent learning, and its normal function is essential for adaptive updating of stimulus-reward associations when rewards are not obtained.

### Amygdala-vlPFC circuits for contingent stimulus-reward learning

Neuroimaging^4^, lesion^1,2^, focused ultrasound^3^, and neurophysiology^5,31,32^ studies in humans and macaques have shown that both amygdala and vlPFC are essential for probabilistic learning and decision-making. Here we found that causally inhibiting the feedforward pathway from amygdala to vlPFC using a chemogenetic approach impaired contingent learning and had a specific effect on how unrewarded trials influenced subsequent behavior. Such a specific role for this pathway is aligned with the findings of a prior neuroimaging study in macaques where Chau, Rushworth, and colleagues reported that functional connectivity between vlPFC and amygdala was increased on lose-shift relative to win-stay trials^4^. Why this pathway appears to be specialized for this function is unclear, but there are several possible explanations. The first is that reward omission may be a more salient or informative cue for guiding learning than receiving a reward. This is because when one option is highly likely to lead to reward, not receiving a reward is a strong indicator that a subject is choosing the option associated with a lower probability of reward and should switch to the alternative. Such a view is, however, not consistent with our finding that the learning rate parameters for unrewarded trials were on average lower than or roughly equal to those for rewarded trials irrespective of whether DCZ was administered or not (**Figure 3C**).

Another possibility is that inhibiting the pathway from amygdala to vlPFC impairs the computation of reward history for each of the choice options. This would appear to fit with 1) the finding that inhibition of this pathway alters reward-outcome representations as opposed to maintaining a representation of the stimulus choice and 2) prior studies that have found that vlPFC is essential for computing reward history in probabilistic settings^2,3^. In the absence of such a temporally extended or model-based learning process, a more myopic or model-free strategy that depends on the most recent outcome would guide behavior. While this strategy would have little effect on win-stay trials, it would have an outsized impact on lose-shift behavior, as animals do not have a model of the task that would cause them to stay with the option that has an overall higher likelihood of leading to reward. Thus, such a model-free strategy based on the outcome of the previous trial would make subjects more likely to switch following an unrewarded trial even when valuations have stabilized later in a session. Notably, such model-free processes are more associated with amygdala-subcortical projections such as those to the ventral striatum, as opposed to amygdala cortical projections^33,34^. Thus, inhibition of amygdala-vlPFC projections may lead to amygdala-subcortical connections guiding learning.

Our task was specifically designed to test whether projections from amygdala to vlPFC contributed to either maintaining a memory trace of the prior choice that was made or signaling the outcome of a choice. We found that this amygdala-vlPFC projections were less involved in this aspect of contingent learning, confirming some prior observations^20^. If amygdala centered circuits are not involved in maintaining information across trace intervals, then which structures are involved? Prior work in humans performing probabilistic learning tasks with trace intervals has highlighted the role of hippocampus^17,18,35^ and interaction between hippocampus and parts of vlPFC^36^. This builds on an extensive literature in lagomorphs and rodents that has pointed to a critical role for the hippocampus^37,38^ as well as medial frontal cortex^22,39^ for maintaining information over periods of time where predictive stimuli are not present. As such, it may be that the amygdala-vlPFC circuit is important for iteratively updating reward history for stimulus reward-associations, whereas hippocampal-frontal circuits are essential for enabling temporally disparate pieces of information to be bound together during learning.

### Interpretational limitations

While we found a clear effect of inhibition of the pathway from amygdala to vlPFC on how unrewarded trials were integrated into current stimulus-reward associations (**Figure 3**), it is important to consider whether this specific effect was caused by our task design. One possibility is that we only saw differences following unrewarded choices because ceiling effects masked alterations in win-stay behavior. Another potential influence of our task design relates to the fact that we did not present an explicit stimulus to subjects when reward omission occurred. By contrast, on rewarded trials the delivery of a food pellet serves as an explicit outcome cue. As such, it is possible that the absence of such a reward omission signal biased how subjects interpreted not receiving a reward, and that this drove the effects on choice behavior that we saw. We note, however, neither of these potential issues can fully account for our findings. First, the effect of pathway specific inhibition was seen for both conditions, even when a trace interval was present for which performance was not at ceiling (**Figure 2**). Second, our findings also appear to match those from Chau, Rushworth, and colleagues, who reported that fMRI functional connectivity between amygdala and vlPFC in macaques was increased on lose-shift trials in a three-choice probabilistic reversal task^4^. Because there was a reversal in their task, subjects had to dynamically sample options and were less likely to choose the best option on every trial and were thus not at the performance ceiling. Taken together, this argues that projections from amygdala to vlPFC appear to be preferentially involved in guiding behavior on trials where a history of reward has to be considered before acting.

Another interpretational issue relates to whether our “pathway specific” manipulation only inhibited a single projection pathway from amygdala to vlPFC. It is becoming increasingly clear that the connections of single neurons in the macaque brain, including those in amygdala, do not target a single area but instead project to multiple locations^40,41^. While we used an intersectional virus approach to target amygdala projections to vlPFC, it is reasonable to assume that these neurons also target other parts of the brain via axon collaterals and that chemogenetic inhibition may have influenced processing in these areas as well. While this feature of brain connectivity and our viral strategy does not alter our conclusions that amygdala projections to vlPFC are involved in contingent learning, it means that a wider network of areas may be involved in such computations, a point we consider below.

### Limbic-frontal networks for learning and foraging

Mapping the projections of individual neurons in the macaque basal and accessory basal nucleus of the amygdala to the frontal cortex has revealed a set of neurons that project to both the vlPFC and dorsal anterior cingulate cortex (dACC)^41^. Notably, this pattern of single neuron projections is not seen in mice^42^, likely because vlPFC, especially area 12, is a primate specialization that appeared during the course of anthropoid primate evolution^43–45^. In addition to connections from amygdala to both of these parts of frontal cortex, macaque vlPFC and dACC are interconnected^46,47^, and interaction between vlPFC and dACC is apparent during reward-guided foraging^3,31,48,49^. Based on this work, it has been hypothesized that while vlPFC tracks the likelihood of receiving a reward so that the best option can be identified, dACC represents the value of alternative options to drive exploration of better alternatives^49–52^. That inhibition of connections from amygdala to vlPFC increased lose-shift behaviors (**Figure 3**) potentially indicates that in addition to reducing the ability of subjects to compute the history of reinforcement for each option, it may have also increased the value of alternative options through interaction with dACC. Thus, single neuron projections from amygdala to vlPFC and dACC may serve to update competing representations that are used to adaptively guide the explore-exploit trade off during foraging. Such a pattern of interactions would be essential for contingent learning in the service of optimal foraging.

## MATERIALS AND METHODS

### Subjects

Three adult male rhesus monkeys (M. mulatta), denoted Cy, He, and Wo, served as subjects. Monkeys weighed between 5.0 and 11.0kg throughout testing, and all were at least 5 years old at the start of testing. Animals were group or pair-housed, although not all within the same group. The largest group consisted of six male macaques; the smallest group was a pair of two. All animals were kept on a 12-hour light/dark cycle, maintained on primate chow and fruit, and had access to ad libitum water. When on a break from study or on weekends each animal was isolated from the group when given food rations in the home cage and regrouped when finished. During behavioral testing, each monkey’s daily food ration was delivered in the test box at the conclusion of behavioral testing. The animals received daily enrichment in the form of a variety of toys that were switched out daily, time in large play cages with hanging enrichment, and were supplemented with fruit and forage mix in their home enclosures. All procedures were reviewed and approved by Icahn School of Medicine Animal Care and Use Committee.

### Surgery

In a dedicated operating room and under full anesthesia, an aseptic neurosurgical procedure to target bilateral virus injections to vlPFC and amygdala was conducted in each macaque. Surgeries were guided by prior T1-weighted structural MRI imaging using the tooth marker method^53^. Anesthesia was induced with ketamine (10mg/kg) and maintained under isoflurane (1-3% for effect). After the skin was opened and the muscles retracted, a large bilateral bone flap was raised over the region of the prefrontal cortex and extending back to around bregma to allow access to the amygdala. A dura flap was then reflected toward the orbit to allow access to the orbital surface in both hemispheres.

Our aim for the surgery was to express inhibitory DREADD receptors into the neurons projecting from amygdala to the vlPFC (**Figure 1C**). To do this, we used an intersectional virus approach where a retrograde virus, canine associated virus-2 expressing cre-recombinase (CAV2-cre) was infused into the vlPFC and a cre-dependent virus expressing the hM4Di-inhibitory DREADD was infused into amygdala, specifically the basal and accessory basal nuclei. Surgery was carried out in two stages; first stereotactic injections were made into the amygdala, and then handheld injections were made into the vlPFC. For the amygdala, injection locations were determined based on pre-operative MRIs to target the basal and accessory basal nuclei of the amygdala. At each location, 1 µl of AAV1-hSyn-HA-DIO-hM4Di (1.78 x 10^12^ vg/ml, Vector Core, National institute of Drug Abuse) was injected at a rate of 0.2 µl per minute. Each animal received 10-12 1ul injections into amygdala in three separate tracks. At the end of each injection the needle was left in place for a further 3 minutes to allow the virus to diffuse away from the injection site. Injections were made in three tracks, two in the middle amygdala and one in the anterior amygdala. At the end of injections in a track, the needle was left in place for 5 minutes to allow for any pressure buildup to dissipate. Individual injections were spaced at least 2mm apart.

For the vlPFC, 25 injections were made into the cortex corresponding to Walker’s areas 12 and lateral agranular insula cortex in each hemisphere. The lateral boundary of the lesion was the convexity of the lateral frontal cortex, and the medial boundary was the fundus of the lateral orbital sulcus. On the inferior frontal convexity, the rostral boundary of the lesion was the mid-point of the principal sulcus, and the caudal boundary was 2 mm posterior to the end of the lateral orbital sulcus. At each location 1 µl of CAV2-Cre (11 x 10^12^ pp/ml, Plateforme de Vectorologique de Montpellier, France) was injected and the needle was held in place for a further 5 seconds to allow the virus to diffuse into the surrounding tissue.

At the conclusion of injections, the dura was closed, the bone flap was sutured into place, the muscles sutured back over the bone flap, and the skin was closed. Following surgery monkeys received postoperative analgesics and anti-inflammatory medication in consultation with veterinarians. Each animal was given at least 2 weeks post-surgical recovery time before training and testing resumed.

### Apparatus and materials

All behavioral testing took place in a sound proofed computer-controlled testing cubicle. The apparatus consisted of a touch-sensitive screen (Hope Industrial Systems, 19” Universal Mount Industrial Monitor Revision H, Model No. HIS-UM19) a pellet dispenser (Med Associates, Inc. Pellet Dispenser 190mg, SN: 10608), as well as two additional disc dispensers, one that dispensed M&Ms and another that dispensed peanuts. Each of these feeders were attached to a food cup located below the touchscreen. A metal lunchbox was located to the left of the food cup and was filled with the large food reward that was delivered at the completion of testing. Infrared cameras positioned at different locations within the test cubicle allowed monkeys’ behavior to be observed from a control room during testing. The cubicle remained dark during testing except for the illumination of the touch-sensitive screen.

For the probabilistic learning task, rewards were 190mg Purified Non-Human Primate Tablets from TestDiet (product 5TUQ), in 3 separate flavors, banana, grape, and raspberry. Delivery of these was controlled by a pellet dispenser positioned above the screen. For the reinforcer devaluation task, rewards were shelled peanuts and chocolate M&Ms (Mars Inc., USA). The lunchbox contained the animal’s daily portion of wet monkey chow, along with a variety of fruits and treats that were selected for each individual monkey based off their preferences (and could include any of the following: banana, orange, mango, grapes, apple, seeds, dates, cranberries, marshmallows, and nuts). Prior to being placed into the testing cubicles monkeys were shaped to enter a transport cage from their home enclosure. The transport cage consisted of 3 solid metal sides, with the fourth containing horizontally oriented bars that allowed the animal to reach the touchscreen, rewards, and lunchbox.

Behavioral tasks on the touchscreen were controlled using Presentation software (Neurobehavioral systems) which sent the appropriate task related signals to the testing apparatus (including when to release pellets, what stimuli to present, and when to open the lunchbox), and further recorded information such as the location of touch responses and their timing. The visual stimuli presented on the touch screen were multi-colored clipart images that were 128 by 128 pixels that were sourced from various on-line repositories.

### Behavioral tasks

#### Dual-pair trace conditioning probabilistic learning task

In the behavioral task, subjects had to discriminate two pairs of stimuli that were probabilistically associated with reward. Each trial started with the presentation of a green lever stimulus (**Figure 1A**). When macaques touched this lever stimulus, two stimuli were presented on the left and right. Monkeys could then choose either of these by touching it, and the chosen stimulus then stayed on the screen for a further 0.5 seconds before a trace interval began and reward feedback was delivered. An ITI then started after reward was delivered or not. To ensure the reward rate for each pair was the same across the course of the session, the ITI was adjusted such that the time from chosen stimulus off to the start of the next trail was always 4 seconds. The probabilities of rewards for the stimuli in each of the pairs were established separately, were independent of one another, and were set at 0.8 and 0.2. One option in each pair was randomly designated as having a reward probability of 0.8 while the other was designated as having a reward probability of 0.2. For one of the pairs of stimuli, feedback (reward or no reward) was delivered immediately after the chosen stimulus was removed from the screen. For the other pair of stimuli there was a 1 second trace interval between the chosen stimulus being removed from the screen and the reward feedback being delivered. Each session consisted of 200 trials with 100 trials of each pair. The presentation of each pair across trials was pseudo-randomly assigned and the left-right position of stimuli was also randomized across trials. Animals had received extensive training on the task and were able to discriminate both pairs of stimuli to a performance level of greater than 80% correct within the first 50-60 trials.

#### Dual-pair trace conditioning probabilistic learning task – DREADD activation

Before animals received surgery, they received different doses of the DREADD activating ligand deschloroclozapine (DCZ) at 0.1 and 0.3 mg/kg 30 minutes prior to performing the dual-pair probabilistic learning task. This testing was conducted to establish whether DCZ alone had any impact on the performance of monkeys Wo, Cy, or He of the dual pair task. Part of these data have been published previously^54^. We conducted 3 sessions with each concentration of DCZ and compared the performance to sessions where saline injections were administered. The DCZ injection days were compared with a vehicle injection day (see section Drug Preparation for more details).

Following virus injections into the brain we then commenced a weekly testing schedule where monkeys were tested 5 days a week as follows. On Monday and Tuesday, we established the monkey’s baseline performance and then on Wednesday and Friday they were administered a DCZ or vehicle injection 30 minutes before behavioral testing. Thursdays served as another baseline/washout day if DCZ was administered on Wednesday. Each monkey completed 10 sessions of testing for each condition, for a total of 20 sessions.

### Reinforcer devaluation task

At the conclusion of probabilistic learning testing, subjects were tested on a reinforcer devaluation task that was administered in the same touch screen apparatus as the probabilistic learning task. Here we used the same automated behavioral task design as described previously^55^. This task has been previously validated in control macaques and shown to be sensitive to lesions of distinct parts of frontal cortex and amygdala^56^. The devaluation task contains several components as detailed below.

#### 60-pair discrimination learning

Monkeys learned to discriminate 60 novel pairs of visual stimuli for rewards. In each pair one stimulus was randomly designated the positive (S+), rewarded stimulus and the other the negative (S-), unrewarded stimulus (**Figure 4A**). For 30 of these pairs, if the monkey chose the positive stimulus, an M&M was delivered. For the other 30 pairs of stimuli, selecting the positive stimulus led to the delivery of a peanut. Once a stimulus was designated positive versus negative their status remained throughout all following training and testing sessions. A key aspect of this task that differs from previous automated versions^55^ is that the lever stimulus used in the probabilistic learning task was also incorporated into the task design. Notably, this feature of the task doesn’t impact the discrimination that the subjects have to perform. Instead, it ensured that there was continuity in the task progression as well as serving as a check on subjects’ motivation to complete the task. In each testing session the pairs of stimuli were presented once in random order, with one stimulus presented on the left of the screen and one on the right. The location of the stimuli on the screen was counterbalanced. Trials began with the presentation of both stimuli and stimuli remained on the screen until the subject made a choice. When one stimulus in a pair was chosen, the unchosen stimulus was removed from the screen and the chosen stimulus remained on for 0.5 seconds before the reward was delivered. There was then a 20 second ITI before the next trial. Reward outcome was deterministic (i.e. selecting the stimulus associated with a reward always resulted in delivery of that reward). Monkeys completed daily sessions of 60 trials until they were discriminating the pairs at a mean of over 80% for 5 consecutive days (i.e. 240 correct out of 300 trials). During these sessions no DCZ or vehicle was administered.

#### Probe tests

Once subjects had attained the performance criteria on the discrimination task, they then started probe and sensory-specific selective satiation tests (**Figure 4B**). Here, subjects’ choices were assessed under two conditions: after either M&Ms or peanuts were devalued using a sensory-specific selective satiation procedure^29^ and in baseline conditions. On separate days we conducted four test sessions, each consisting of 30 trials where only the positive, reward associated stimuli were used. On each trial, stimuli associated with M&Ms and peanuts were presented together for choice, with the constraint that an M&M associated stimulus was always paired with a stimulus that was associated with peanuts. The stimulus pairs were generated pseudorandomly for each session with the constraint that monkeys were only ever presented with unique combinations of stimuli once. The testing week for this experiment consisted of 4 behavioral days, and sessions were counterbalanced so that there were always 3 days between sessions where the selective satiation procedure was conducted and subsequent training. Practically, each week monkeys completed the 60 pair discrimination task on Monday and Wednesday. These sessions were used to confirm that subjects knew the stimulus reward associations and to maintain a high level of performance. Baseline probe tests that were not preceded by the selective satiation procedure were conducted on Tuesdays, while probe tests that were preceded by the selective satiation procedure occurred on Thursdays. Establishing the testing progression this way allowed us to treat Friday-Sunday as drug/satiation washout days where monkeys were not tested.

#### Sensory-specific selective satiation

On days where the selective satiation procedure was conducted, approximately 24hr after the last feeding, animals were provided with 300 grams of M&Ms or peanuts in their home cage. The monkey was allowed to eat the food while being discreetly observed through a door viewer until they stopped actively eating, or for 20 minutes. After 20 minutes uneaten food was removed from the home cage and later measured to determine the amount consumed. Animals were then given an additional 5-10 minutes to empty their cheek pouches. The selective satiation procedure usually started 30 minutes prior to the start of the probe task. Each subject completed at least two sessions where vehicle or DCZ (i.m. 0.1 mg/kg) were administered 5 minutes before the start of the 20-minute selective satiation procedure, and 30 minutes before the start of the behavioral task.

### Drug preparation

Both vehicle and DCZ were prepared using sterile procedures and previously published drug preparation approaches for behavioral testing in macaques^25,57^. The vehicle consisted of 0.5% Acetic acid, 50% 1 M Sodium Acetate, and 49.5% 0.2 M Sodium hydroxide (NaOH). DCZ (Medchemexpress, Monmouth Junction, NJ) was stored at 4° C in a designated fridge, while the acetic acid, sodium acetate, and NaOH was stored at room temperature (all obtained from Fisher Scientific). Each solution of DCZ or vehicle ultimately had a pH between 5.5 and 6.0. When DCZ was administered, the drug was first dissolved in acetic acid and sodium acetate and then diluted with NaOH. The vehicle remained consistent across both drug doses (0.1mg/kg, or 0.3mg/kg of DCZ). However, the concentration of each dose was altered so the total volume of each injection was the same no matter the dose, e.g. a 0.1mg/kg dose had a concentration of 1 mg/mL, while a 0.3 mg/kg dose had a concentration of 3 mg/mL. DCZ or vehicle solution was prepared fresh daily and within 30 min of usage.

### Statistical analysis

Statistical analyses were conducted in MATLAB version 2024a. For the probabilistic learning task, we analyzed subjects’ higher probabilistic option choices (0 or 1), initiation times and reaction times. Initiation time was defined as the amount of time it took subjects to press the green lever stimulus after it was presented. Reaction time was defined as the amount of time from the green lever stimulus being touched to one of the subsequently presented pair of stimuli being touched. We used repeated measures ANOVAN to analyze the initiation and reaction times. Specifically, using the “anovan” function in MATLAB we included main effects of monkey, pair, and drug with inactivity, and reaction time as dependent variables. Session was included as a random effect. To compare the performance of different the subjects on the probabilistic learning task, we conducted a binomial logistic regression using the fitglme (fit generalized linear mixed effects model [GLME]) function in Matlab. This analysis included main effects for monkey (3 levels), pair (2 levels), block (2 levels), and drug (2 levels). As before, session (10 levels) was included in the model as a random effect.

For the reinforcer devaluation task, we computed a difference score between the baseline sessions and those preceded by the selective satiation procedure to reduce the value of stimuli associated with peanuts or M&Ms^29^. To control for local variation in preferences between peanuts and M&Ms, the total number of times the monkeys chose the option associated with the M&M stimuli in baseline sessions was subtracted from the total in session preceded by selective satiation in the same week. Difference scores were analyzed using repeated measures ANOVA with main effects of monkey (3 levels) and drug (2 levels).

### Reinforcement learning model

To estimate the influence of rewarded and unrewarded trials on subjects’ behavior, we fitted a three-parameter reinforcement learning model to the choice data. This model was fitted separately for each pair of stimuli in a session and produced estimates of stimulus value and choice probability, for each stimulus on each trial. A learning rate for rewarded trials, a learning rate for unrewarded trials, and an inverse temperature parameter was also computed for each session. The model updates the value, *v*, of a chosen option, *i*, based on reward feedback, *r* in trial *t* as follows:

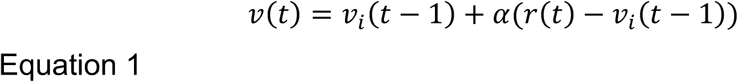

Thus, the updated value of an option is given by its old value, *v_i_(t – 1)* plus a change based on the reward prediction error *(r(t) – v_i_(t – 1))*, multiplied by the learning rate parameter, *α* that is separated for the rewarded (*α*_rewarded_) and unrewarded trials (*α*_unrewarded_). At the beginning of each session, the value, v, of both stimuli is set to 0.5. The three free parameters (the learning rate parameters, *α*, and the inverse temperature, *β*, which estimates how consistently animals choose the highest valued option), were fit by maximizing the likelihood of the choice behavior of the monkeys, given the model parameters. Specifically, we calculated the choice probability *d_i_(t)* using the following:

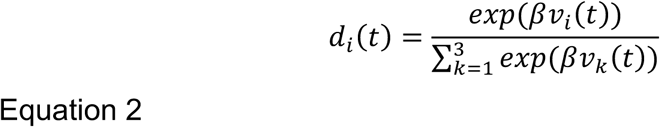

And then calculated the log-likelihood as follows:

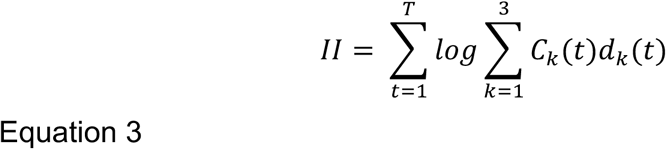

Where *c_k_(t) = 1* when the subject chooses option *k* in trial *t* and *c_k_(t)* = 0 for all unchosen options, meaning that the model maximizes the choice probability (*d_k_(t)*) of the actual choices the monkeys made. *T* is the total number of trials that monkeys completed in a session. Parameters were maximized using standard techniques^58^. Learning rate parameters were drawn from a normal distribution with a mean of 0.5 and a standard deviation of 3. The inverse temperature parameter was drawn from a normal distribution with a mean of 1 and a standard deviation of 5. These distributions were chosen because learning rates in probabilistic settings should be considerably less than 1, given the stochastic nature of reward delivery, and positive inverse temperatures indicate that choices are biased towards higher reward values. Model fits were repeated 1000 times to avoid local minima. To ensure that the estimated parameters stayed within appropriate bounds we set a min-max value of 0 and 1 and 0 and 30 for the learning rate and inverse temperature parameters respectively. Any sets of choices where the parameters failed to converge after 1000 iterations or that could not be fitted using the model were excluded from further analysis. This happened in 5 out of the 120 sessions pairs available for analysis. Free parameters from the reinforcement learning model were compared using ANOVA with main effects of subject (3 levels), pair (2 levels), and drug (2 levels).

### Histological processing

Perfusion and histological processing was performed similarly to our previous studies^59^. Animals were deeply anesthetized with a sodium pentabarbitol solution and perfused transcardially with 0.1M phosphate buffered saline (PBS) and 1% (w/v) formaldehyde solution derived from depolymerized paraformaldehyde followed by 0.1M PBS and 4% (w/v) formaldehyde solution derived from depolymerized paraformaldehyde. Brains were removed and immersed in this same formaldehyde solution overnight. Following post-fixation, brains were transferred to a solution of 2% DMSO and 10% glycerol in 0.1M PBS for 24–48 h, then transferred to a solution of 2% DMSO and 20% glycerol in 0.1M PBS for at least 24 h. Cryoprotected brains were blocked in the coronal plane, then flash frozen in −80°C isopentane. After flash freezing, the brains were removed, dried, and stored at −80°C until sectioning. Tissue was cut serially in 50 µm coronal sections on a sliding microtome (Leica SM 2010R) equipped with a freezing stage.

To confirm DREADD expression, sections through the amygdala and the vlPFC were selected and washed with PBS solution containing 0.3% Triton X-100 (TX-100). Endogenous peroxidase was quenched by incubation in 0.6% hydrogen peroxide in PBS for ten min. The sections were washed, then blocked for 1 h in a solution of 10% bovine serum albumin (Cat #BP671-1, Thermo Fisher Scientific, Waltham, MA), 10% normal goat serum (Cat #PI31873, Invitrogen, Waltham, MA), and 0.3% TX-100 in PBS. Primary antibody against the hemagglutinin tag (raised in rabbit, clone C29F4, 1:800, Cell Signaling Technology, Danvers, MA; Cat #3724, RRID:AB_1549585) was added to the blocking solution and sections were incubated overnight. Incubation in primary antibody was followed by washes and a 2-h incubation in anti-rabbit biotinylated secondary antibodies (1:200, Vector Laboratories, Burlingame, CA, RRID:AB_2313606). Sections were then washed and incubated for at least 1.5 h in avidin-biotin complex solution (1:200; Vectastain standard kit, Vector Laboratories, RRID:AB_2336819). Finally, sections were washed, incubated in a 3-3’-diaminobenzidine tetrahydrochloride (DAB) solution for 10 min, and 0.006% hydrogen peroxide was added to the solution. Sections were carefully observed for a reaction and were placed in buffer solution after approximately 4 min, when staining was visually apparent. Sections were mounted on glass slides, air dried, dehydrated in ascending concentrations of ethanol, and cleared in d-limonene, then coverslipped with DPX mounting medium.

Images of sections were acquired using a Zeiss Apotome 2 microscope equipped with Q-Imaging and Hamamatsu digital cameras, a motorized stage, and Stereo Investigator software (MBF Bioscience; RRID:SCR_002526). Images were acquired using consistent exposure time and brightness settings within the same section. Any adjustments made afterward (e.g., to increase brightness) were applied uniformly to the image. To obtain wide field images of the entire section, the outline of the tissue was defined using a 5x lens. Tiled images of a single section were obtained sequentially, constrained by the boundaries of the tissue, and a 10% overlap and image stitching was used to create a composite image of all tiled photos.

## AUTHORS CONTRIBUTIONS

Conceptualization and Methodology: JMF, PHR; Investigation, Data curation, Analysis, Software and Visualization: JMF, CE, JEB, PHR; Writing – Original Draft and Review/Editing: JMF, CE, JEB, PHR; Funding Acquisition and Supervision: PHR.

## ACKNOWLEDGEMENTS

This work was supported by a National Institute of Mental Health BRAINS award to PHR (R01 MH110822), a young investigator grant from the Brain and Behavior Foundation (NARSAD) to PHR, and seed funds from the Icahn School of Medicine at Mount Sinai to PHR. We would like to thank Chris Richie at NIDA for providing the AAV CRE-dependent viruses and Mark Baxter for providing access to testing equipment.

## SUPPLEMENTARY INFORMATION

**Figure S1:**
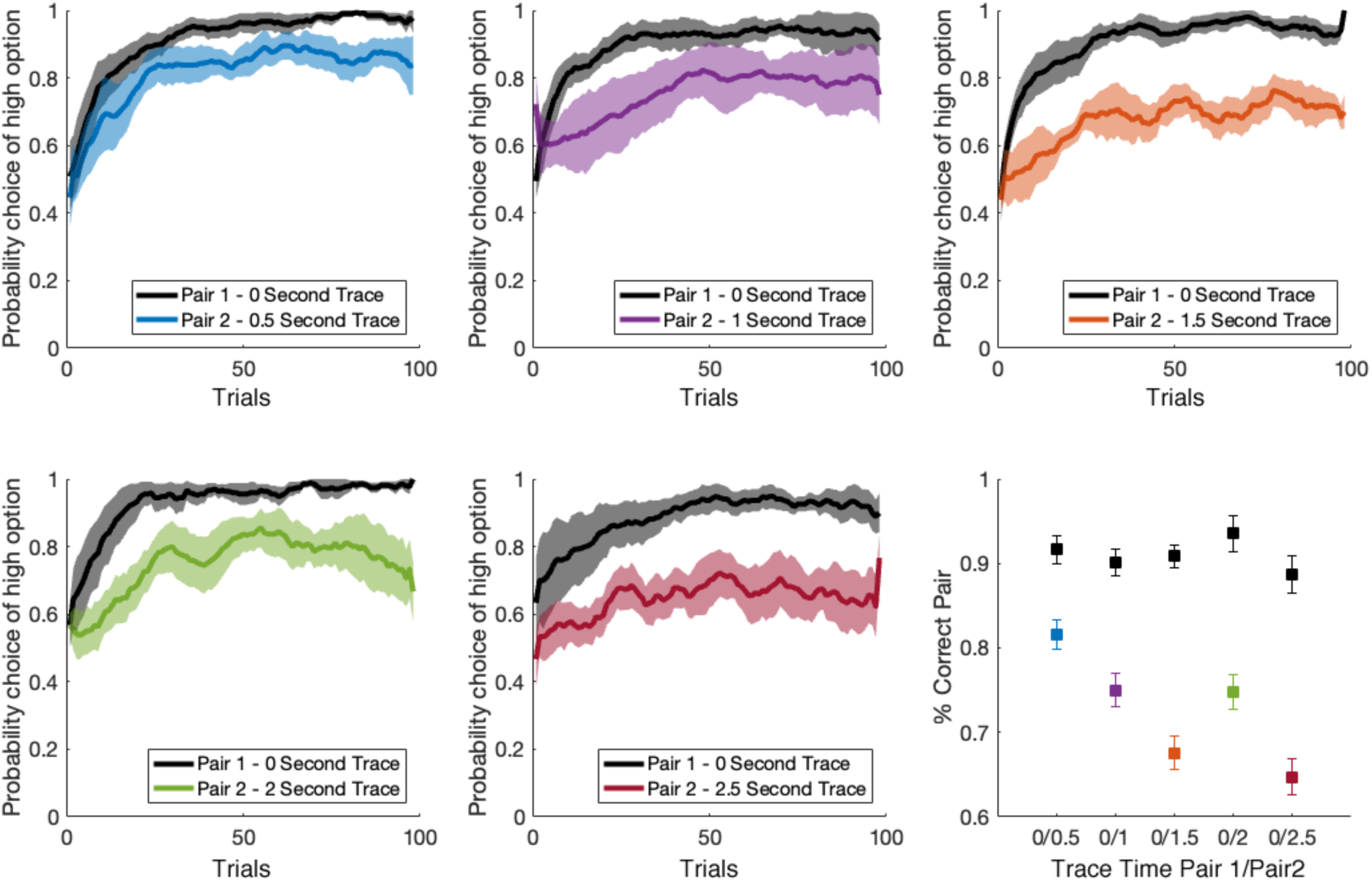
Probabilistic learning performance with trace intervals of different lengths in the two-option probabilistic learning task. Mean (+/- SEM) high reward option choice for a pair of stimuli where reward is delivered immediately (0 second trace, black line) and a pair of stimuli where reward is delivered after a trace interval of length 0.5 – 2.5 second. Performance is from a group of 4 macaques trained to perform the where in each pair one stimulus was associated with 0.8 probability of reward, whereas the other was associated with 0.2. The general task progression is the same as in Figure 1. Mean (+/- SEM) choice of high reward option for each pair of stimuli is shown in bottom right corner.

